# Isolation of functional supramolecular attack particles (SMAPs)

**DOI:** 10.1101/2025.11.10.687675

**Authors:** Ashwin Jainarayanan, Vineethkrishna Chandrasekar, Vinnycius Pereira Almeida, Marcus A. Widdess, Claire C. Staton, Agathe Bernand, Veronika Pfannenstill, Sarah Flannery, Helena Watson, Amanda Wicki, Orthi Onupom, Mirudula Elanchezhian, Jesusa Capera, Lina Chen, Salvatore Valvo, Nithishwer Mouroug Anand, Ranjeet Singh Mahla, Elke Kurz, Camille Franchet, Florence Dalenc, Loïc Ysebaert, Amandine Michelet, Dhanu Gupta, James H. Naismith, Lynn B. Dustin, Marco Fritzsche, Roman Fischer, Salvatore Valitutti, Carol S. Leung, Matthew J. A. Wood, Michael L. Dustin

**Affiliations:** Kennedy Institute of Rheumatology, Nuffield Department of Orthopaedics, Rheumatology and Musculoskeletal Sciences, University of Oxford, Oxford, UK; Institute of Developmental and Regenerative Medicine, Department of Paediatrics, University of Oxford, Oxford, UK; Ludwig Cancer Research, Nuffield Department of Medicine, University of Oxford, Oxford, UK; Institut National de la Santé et de la Recherche Médicale (INSERM) UMR1037, Centre de Recherche en Cancérologie de Toulouse (CRCT), Toulouse, FR; Discovery Proteomics Facility, Nuffield Department of Medicine, University of Oxford, Oxford UK; Rosalind Franklin Institute, Harwell Campus, Oxford, UK; Department of Biology, University of Oxford, Oxford, UK; Department of Biochemistry, University of Oxford, Oxford, UK; Department of Pathology, Institut Universitaire du Cancer-Oncopole de Toulouse, Toulouse, FR; Department of Medical Oncology, Institut Universitaire du Cancer-Oncopole de Toulouse, Toulouse, FR; Department of Hematology, Institut Universitaire du Cancer-Oncopole de Toulouse, Toulouse, FR; Division of Structural Biology, Nuffield Department of Medicine, University of Oxford, UK; Centre for Immuno-Oncology, Nuffield Department of Medicine, University of Oxford, UK

**Keywords:** cytotoxicity, particles, glycoproteins, immuno-oncology, nanotechnology

## Abstract

High density cultures of the GMP compatible NK-92 natural killer cell line release heterogeneous extracellular particles (EP), notably extracellular vesicles (EV) and non-vesicular extracellular particles (NVEP). The NVEP include a small proportion of supramolecular attack particles (SMAPs), the direct cytotoxic potential of which suggests therapeutic promise, yet scalable enrichment and functional evaluation alongside other extracellular particles (EPs) have been lacking. Here, we develop a high-resolution serial size-exclusion liquid chromatography (SELC) workflow that resolves EPs into 4 fractions F1-F4, in which EV are enriched in F2 and SMAPs are enriched in F3. Multi-modal characterization using Nanoparticle Tracking Analysis (NTA), Nano Flow Cytometry (Nano FCM), Transmission Electron Microscopy (TEM), Total Internal Reflection Microscopy (TIRFM) and Proteomics) demonstrated that F2 NK-92 EVs are ∼130–150 nm particles enriched for complement proteins, whereas SMAPs (F3) are 100–110 nm particles enriched for cytotoxic proteins (PRF1, GZMB) and SMAP shell components (e.g., THBS1, THBS4). Functionally, Ca^2+^-stabilized SMAPs (F3) trigger caspase-3–dependent apoptosis in tumor cell lines with robust dose responses, while F2 lacks direct cytotoxicity in vitro. Ex vivo, F3 SMAPs kill patient-derived breast cancer and chronic lymphocytic leukaemia cells (CLL) in a concentration-dependent manner. In NSG mice, intra-tumoral treatment with SMAPs (F3) restrains the growth of aggressive B16F10 melanoma and PANC-1 pancreatic cancer, confirming in vivo functionality. These results establish a scalable method to isolate and compare NK-derived particle classes and provide a foundation for targeted SMAP engineering as a promising approach for treatment of solid tumours.

**Significance Statement:** Cell therapies face barriers in solid tumours, including lack of entry and suppression. NK cells release extracellular particles (EPs), including supramolecular attack particles (SMAPs), that can traffic into tumour tissue. We define a practical, scalable chromatography process that resolves NK-derived SMAPs from other EPs and demonstrate that SMAPs induce caspase mediated apoptosis *in vitro* and engage in cytotoxic lymphocyte independent anti-tumour activity *in vivo*. These insights open a route to engineering and testing SMAP-based therapeutics to overcome limitations of whole-cell approaches in solid cancers.

## Introduction

Cytotoxic lymphocytes—natural killer (NK) cells and effector CD8⁺ T cells—eliminate malignant targets through granule exocytosis (perforin and granzymes) and death-receptor pathways, and they have inspired cell therapies that achieve durable remissions in select hematologic malignancies. The solid-tumor setting remains challenging due to antigen heterogeneity, immunosuppressive tumor microenvironments, and poor persistence and trafficking of infused cells. These constraints motivate cell-free approaches that retain cytolytic potential while improving tissue penetration, dosing control, and manufacturing consistency^1^.

Immune cells release a spectrum of extracellular particles (EPs), including membrane-bounded extracellular vesicles (EVs) and non-vesicular extracellular particles (NVEPs). EVs have been associated with cytotoxic proteins and immune ligands with immunomodulatory potential and have been investigated as antitumor agents and delivery vehicles. A distinct class of NVEPs—supramolecular attack particles (SMAPs)—are ∼100–120 nm proteinaceous particles with a thrombospondin-rich shell that encapsulates perforin and granzymes, enabling autonomous killing independent of effector cells.

These particles are known to be secreted by both cytotoxic T cells and NK cells^2,3^. Mechanistic understanding regarding SMAP composition and biogenesis are being refined, thrombospondin-4 cooperates with thrombospondin-1 to build the shell, and multicore granules have been identified as a source of SMAP cargo^4,5^.

Further mechanistic and functional dissection of SMAP function has been impeded by their rarity within pooled EPs and by the difficulty of separating vesicular from non-vesicular particles at preparative scale, for example in in vivo studies, without altering function. The latest ISEV consensus (MISEV2023) emphasizes multimodal readouts and cautions that EV preparations often contain multiple particle types, including non-vesicular entities^6^. Proteomic comparisons highlight EV subtype heterogeneity^7^, and size-exclusion chromatography (SEC) is widely used to improve particle-level resolution^8,9,10^. In addition, EV preparations frequently display a surface “protein corona” acquired from biofluids; corona composition can recruit complement components, modulate hydrodynamic size, and influence biodistribution and immunogenicity^11,12,13^. Accordingly, clean resolution and functional testing of a discrete particle class are essential to link its respective composition with mechanism.

Here we establish a serial, high-resolution size-exclusion liquid chromatography (SELC) workflow using the GMP-compatible NK-92 line to resolve functionally distinct particle populations from culture supernatants. The workflow yields a fraction enriched for EV-associated markers and complement proteins—referred to here as CompEVs (for “complement-EVs”)—and a fraction enriched for SMAP components (perforin, granzyme B, thrombospondin-1) referred to as SMAPs. Orthogonal sizing and imaging, immunoblotting, and quantitative proteomics define compositional differences between these fractions, and functional assays map these differences to distinct antitumor effects *in vitro*, *ex vivo*, and *in vivo*.

## Results

High density NK-92 cells cultures released cytotoxic EPs (purified by PEG precipitation, PEG-EPs) by 96 hours, but not by 48 hours (Fig 1A). Based on immunoblotting, the 96 hours contained EVs (CD63, CD81 and TSG101 positive) and SMAPs (THBS1 and GZMB positive), and largely lacked contaminating cellular membrane and organelles (GMP120, Calnexin, Cytochrome C and Actin negative) (Fig 1B). Particle analysis using WGA to track all PEG-EPs, CD81 to identify EVs and PRF1 and GZMB to mark SMAPs, revealed that 4% of PEG-EPs were SMAPs with PRF1 and GZMB, and there were GZMB and PRF1 single positive NVEPs (12%) and EVs (12%) present in the total released EPs (Fig 1C). We explored the potential of tandem size exclusion liquid chromatography (SELC) to resolve SMAPs from EV within the EP from NK-92 cells (Fig. 1D). SELC on Sepharose 4 Fast Flow separated an EP-rich void volume, which elutes first, from soluble proteins, which elute later (Fig. 1D, E). Nanopaticle tracking analysis demonstrated significant size heterogeneity in the NK-92 EPs from Sepharose 4 Fast Flow SELC (Fig. 1F). Proteomic profiling of NK-92 EPs (purified by 4FF column, EPs) included the cytotoxic T lymphocyte SMAP signatures obtained directly from synaptic secretions^3^ (Fig S1A), demonstrating that SMAPs can be enriched by SELC. Furthermore, particle analysis by immunostaining revealed PRF1⁺/GZMB⁺ SMAP-like NVEPs and double-positive EVs with CD81 and PRF1 or GZMB (Fig S1B), demonstrating that all populations in the starting EP’s from NK-92 cells were recovered by SELC with an apparent increase in the ratio of NVEP to EV. This EP fraction was then subjected to further high-resolution size exclusion chromatography (HR-SEC) to further fractionate the EPs. Using Sephacryl S-1000 (or Sephacryl S-500), we reproducibly resolved four fractions (F1–F4) with dominant peaks in F2 and F3 (two fractions from Sephacryl S-500 SELC correspond to these fractions) (Fig 1G). Nanoparticle tracking analysis (NTA) assigned diameters of ∼130–150 nm for F2 and ∼100–110 nm for F3 from the Sephacryl S-1000 SELC (Fig 1G,H).

**Fig 1.**
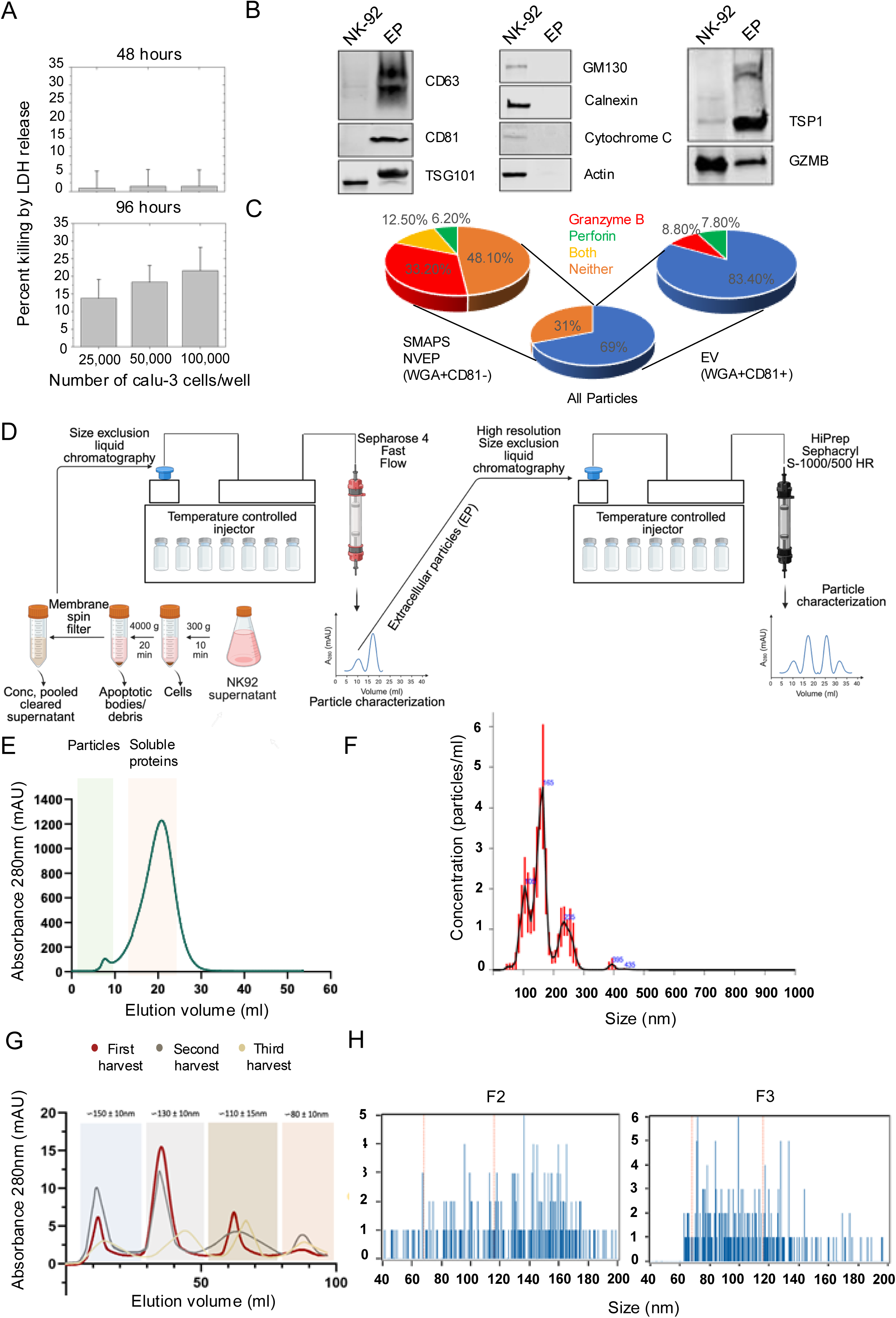
High resolution serial Size Exclusion chromatography enables isolation of distinct population of NK92 secreted particles. (A) PEG precipitated particles from NK92 supernatant were added to Calu-3 cells for 48 and 96 hours and subjected to LDH release assay to measure cytotoxicity (B) NK92 PEG precipitated particles were subject to immunoblotting alongside NK92 cell lysate and probed for EV and SMAP markers labeled in the figure (C) PEG precipitated NK92 particles were adsorbed on to glass coverslips and analyzed for WGA, CD81, PRF1 and GZMB by TIRFM. CD81+ were counted as EV and WGA^+^ CD81^-^ particles were considered NVEPs. The percentage of EV and NVEP that expressed each marker were determined and displayed in the pie chart. (D) Schematic of the isolation and fractionation of secreted particles from NK cell culture supernatant, to separate out secreted particles from soluble proteins using Sepharose 4FF. Extracellular particles from 4FF were further purified using high-resolution size exclusion liquid chromatography (SELC) on Sephacryl S-1000/500 HR columns. A temperature-controlled injector ensured consistent column performance and reproducibility. Distinct fractions were collected based on elution profiles, resulting in separation of particle-rich fractions (F2, F3). (E) Representative SELC chromatogram showing the intensities of the secreted particle peak and the soluble proteins peak (F) Nanoparticle Tracking Analysis (NTA): Concentration (particles/mL) and size distribution profiles of secreted particles across the pooled sample, with a dominant peak at ∼140 nm and additional smaller particle populations. (G) SELC chromatograms: Absorbance at 280 nm (mAU) of sequential harvests of NK-92 culture supernatants (1st–3rd harvests), showing reproducible separation into four distinct fractions. Approximate mean particle diameters were determined by NTA: ∼150 ± 10 nm (Fraction 1), ∼130 ± 10 nm (Fraction 2), ∼110 ± 15 nm (Fraction 3), and ∼80 ± 10 nm (Fraction 4). Fraction 2 consistently yielded the strongest absorbance peak, corresponding to the major particle population. (H) NanoFCM analysis: High-resolution single-particle size distributions of F2 and F3 fractions revealed enrichment of relatively larger F2 particles (∼150 nm), while F3 particles were smaller (∼100 nm).

We next sought to determine if the major fractions F2 or F3 were enriched for EVs or SMAPs. To begin to assess this, the presence of key EV and SMAP proteins was analysed by immunoblotting. To our delight, when equal amounts of protein were loaded, F2 was enriched with EV markers FasL and CD81, whereas F3 was enriched in SMAP markers THBS1, PRF1 and GZMB (Fig 2A). To identify other proteins constituents, we subjected F2 and F3 to mass spectrometry based proteomic analysis. Initially, we normalized the counts to EV marker GAPDH^14^, which allowed us to track EV and determine if we have enriched SMAPs over EV in F3. Surprisingly, the most abundant proteins in F2 were complement proteins. However, we did find EV in F2 under the complement signals with multiple EV markers having a value close to that of GAPDH. Importantly, granzyme mediated killing pathway proteins were enriched over GAPDH in F3, consistent with this fraction not only containing SMAPs, but enriching them over EV. Pathway enrichment analysis confirmed that, F2 was indeed enriched for proteins involved in regulation of complement pathway, whereas F3 was enriched for proteins involved in Granzyme mediated programmed cell death pathways (Fig 2 B-D, Fig S2 A-B).

**Fig 2.**
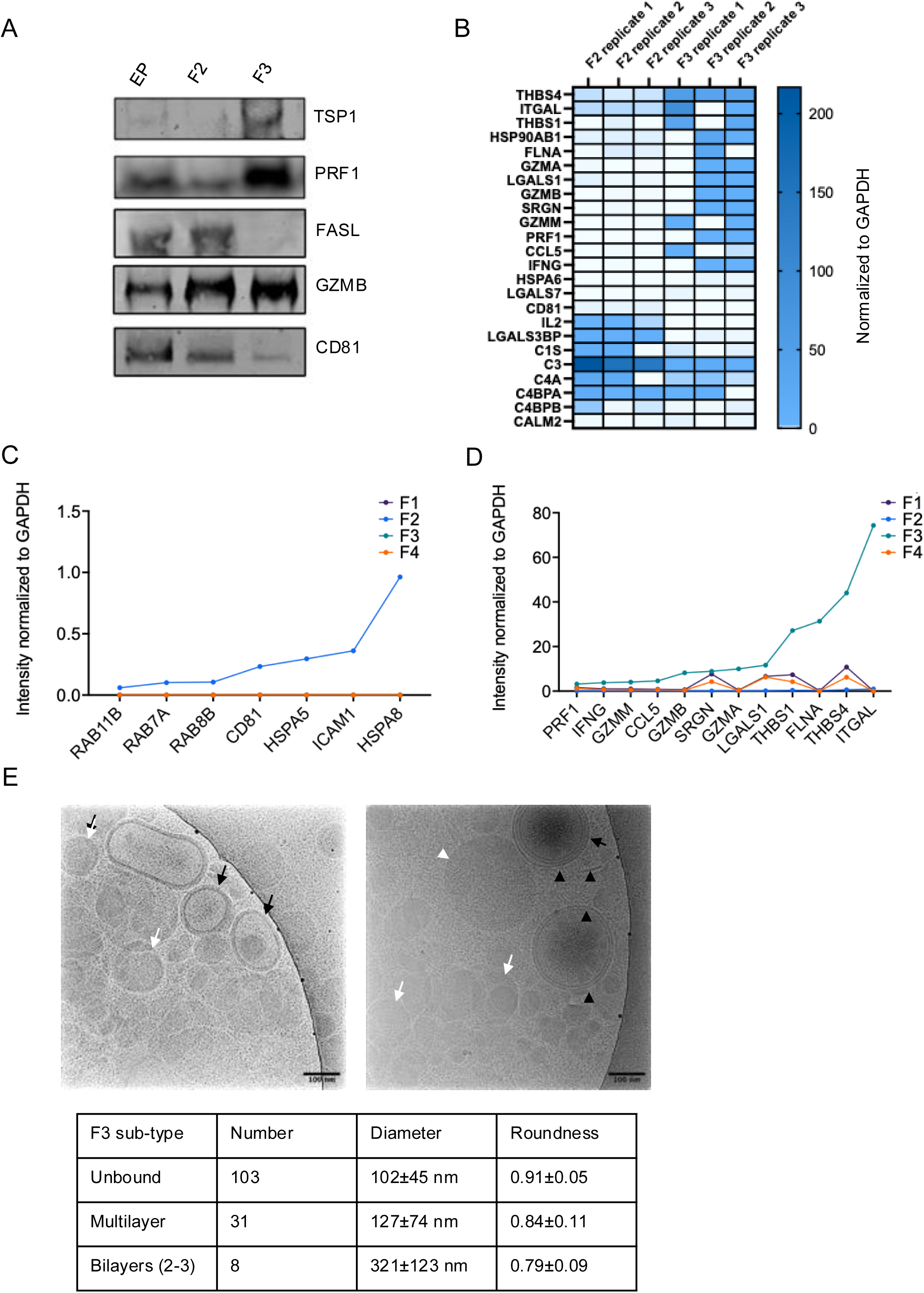
High resolution serial Size Exclusion chromatography enables isolation of SMAPs (F3) and Extracellular Vesicles (EV) (F2) (A) Immunoblotting for SMAP and EV markers across 4FF derived EP particles and S1000/500 derived F2 and F3 particles (B) GAPDH-normalized intensities of SMAP and EV signature proteins along with complement proteins plotted across proteomics replicates (C) & (D) Proteomics analysis of F1-F4, the average GAPDH-normalized intensity of SMAP related markers (D) and Exosome markers (D) across all fractions (F1-F4) is shown as a line graph. Sorted by lowest to highest on EV markers (C) and SMAP markers (D) (E) CryoEM of NK92 particles, representative images and characterization of morphology in a summary table.

Cryo-electron microscopy (Cryo-EM) was performed on F3 to initially assess structural features of the particles. EV have been extensively studied by Cryo-EM^15^, F3 contained heterogeneous particles most of which were unbound (n = 103), although some particles appeared to have 2-3 bilayers (n = 8) or multilayer structures (n = 31). The unbound particles had the greatest roundness were mostly round, and could have nested inclusions of different densities. None of the particles observed looked like classic EVs (Fig 2E).

Thus, F3 from tandem SELC was highly enriched for cytotoxic proteins with a SMAP-like protein signature and morphology and was selected for further functional analysis.

We had previously demonstrated that surface presented SMAPs were cytotoxic^3,5^ but had not had access to SMAPs in suspension. To functionally validate the NK-92 SMAP suspensions, we first performed in vitro cytotoxicity assays, where we measured uptake of live/dead dye by flow cytometry. Our initial results were inconsistent. Parallel analysis of related thrombospondin encapsulated dense particles (ThrEDs) from epithelial cell line A549 suggested that maintaining extracellular Ca^2+^ isolation and storage could be important for integrity of thrombospondin-containing particles^16^, including SMAPs. When we stored SMAPs (F3) with 1 mM Ca^2+^, functional assays were more consistent. F3 stored with Ca²⁺ killed NALM-6 leukemia cells within 24 hours, whereas F3 stored without Ca^2+^ or F2 stored with or without Ca^2+^ did not (Fig. 3A). This result demonstrates direct cytotoxic activity of SMAP suspensions (F3) consistent with prior results on substrate attached SMAPs from T cells^3^. Thrombospondins are Ca²⁺-binding matricellular proteins and, thus, Ca^2+^ may further stabilize the SMAP shell^17^. In addition, Ca²⁺ stabilization preserved SMAP activity for ≥1 month at 4 °C (Fig. S3), supporting practical handling. Also, F3 SMAPs displayed reproducible cytotoxic effects on NALM-6 cells at 72 hours in a dose dependent manner across preparations (Fig. 3B). Furthermore, we observed a time and concentration-dependent activity against JY Lymphoblastoid cell line, validating SMAP activity in multiple targets (Fig. 3C). Mechanistically, live-cell imaging showed caspase-3 activation preceding membrane permeabilization (Fig. 3D–E; Video S1).

**Figure 3.**
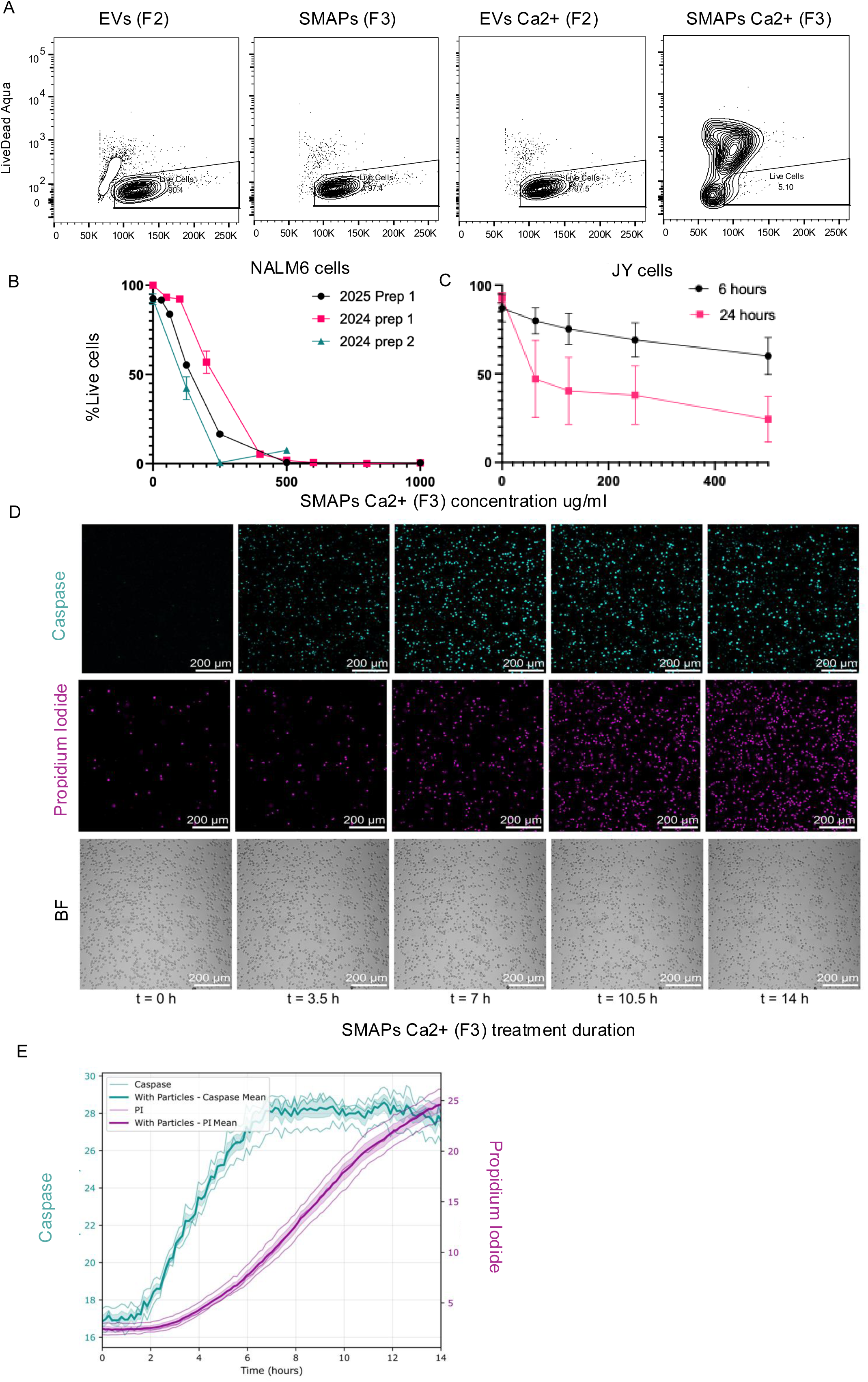
Calcium-dependent stabilization of F3 particles enables SMAP-mediated cytotoxicity through caspase-3 activation, while F2 particles lack cytotoxic activity in vitro. (A) Flow cytometry-based live/dead cell viability assay of NALM-6 acute lymphoblastic leukemia cells following 24-hour treatment with 500 μg/mL of either SMAPs (F3 fraction) or EV (F2 fraction) in the presence or absence of calcium supplementation during storage post fractionation. Results demonstrate calcium-dependent cytotoxicity of F3 particles but not F2 particles. (B) Dose-response cytotoxicity analysis of SMAPs (F3 fraction) against NALM-6 cells measured after 72 hours of treatment, showing reproducibility across independent particle preparations (n=3 biological replicates). (C) Time-course and concentration-dependent cytotoxic effects of F3 SMAPs on JY B-lymphoblast cells assessed by flow cytometry live/dead assay over multiple time points and concentrations (n=4 biological replicates). (D) Representative fluorescence microscopy images from live-cell imaging experiments monitoring caspase-3 activation (green fluorescence) and propidium iodide (PI) uptake (red fluorescence, indicating membrane permeabilization and cell death) over a 16-hour time course following SMAP (F3) treatment. Images demonstrate temporal progression of apoptotic signaling and cell death. *(E)* Quantitative analysis of fluorescence intensity for both PI uptake and caspase-3 activation over the 16-hour live imaging period, showing kinetics of SMAP-induced apoptotic cell death pathway activation.

We extended these experiments to *ex vivo* treatment of patient derived cancer cells to evaluate clinical relevance. Notably, treatment with Ca^2+^-stabilized F3 SMAPs produced a concentration-dependent cytotoxic effect in patient-derived breast cancer (n = 2) and chronic lymphocytic leukemia (CLLs) (n = 3) cells over 24 h (Fig. 4A–C). This ex vivo activity generalizes SMAP killing beyond cell lines and supports translational relevance for diverse tumour types.

**Figure 4.**
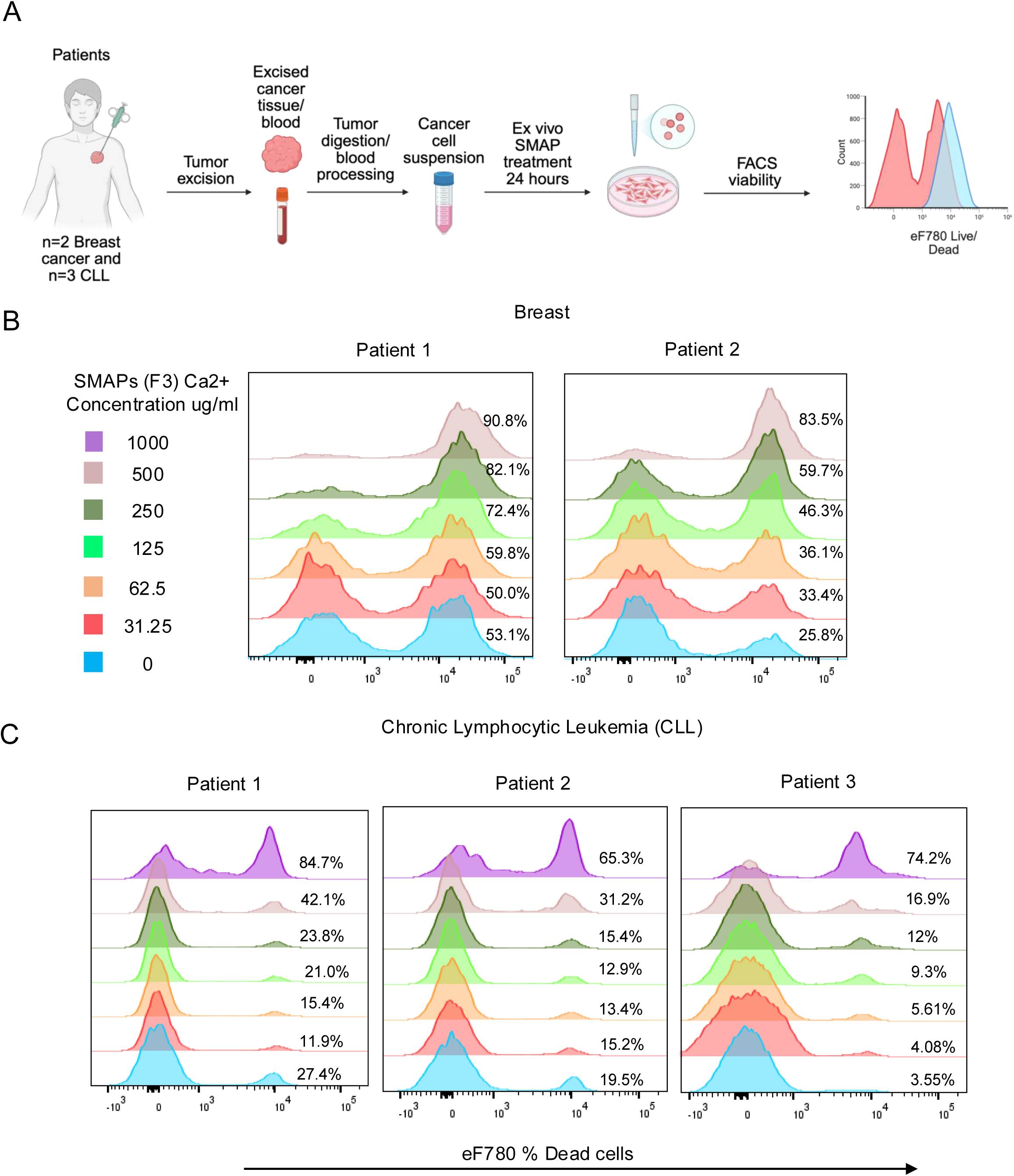
SMAPs (F3) exhibit dose-dependent cytotoxicity against primary tumor cells isolated from breast cancer and chronic lymphocytic leukemia (CLL) patients. (A) Experimental workflow schematic illustrating the processing of patient-derived tumor samples for ex vivo cytotoxicity assessment. Fresh breast cancer surgical specimens were enzymatically and mechanically digested to generate single-cell suspensions. CLL cells were isolated from peripheral blood of patients. Cells were subsequently treated with varying concentrations of SMAPs (F3 fraction) and analyzed by flow cytometry-based viability assays. (B) Dose-response cytotoxicity curves of SMAPs against primary breast cancer cells isolated from two independent patient tumor samples (n=2 patients). Cells were treated with SMAPs at concentrations ranging from 0 to 500 μg/mL for 24 hours, and viability was quantified using flow cytometry live/dead discrimination assays. (C) Dose-response cytotoxicity analysis of SMAPs against primary CLL cells obtained from three independent patient samples (n=3 patients). Viability assessment methods were identical to those described in (B). Data are presented as individual patient responses as indicated. Cell death is noted as % of dead cells relative to untreated control.

To evaluate if Ca^2+^-stabilized F3 SMAPs were functional in vivo, we used NOD scid gamma (NSG) mice, to rule out lymphocyte dependent tumour rejection, and implanted them subcutaneously with an aggressive fast growing B16F10 melanoma line (B16) or a slower growing PANC-1 pancreatic cancer line, a week before treatment. As the NK-92 source had no engineered specificity towards tumour or tissue-specific targets, we intratumorally injected Ca^2+^-stabilized F3 SMAPs five times every two to three days for 2 weeks and tracked mouse weight and tumour growth (Fig 5 A). In NSG mice bearing PANC-1 pancreatic tumours, repeated intratumoral dosing was well tolerated (no weight loss) (Fig. 5B). Ca^2+^- stabilized F3 SMAPs controlled B16F10 and PANC-1 tumour growth significantly (Fig. 5C–E), confirming a lymphocyte independent direct anti-tumour effect in vivo.

**Figure 5.**
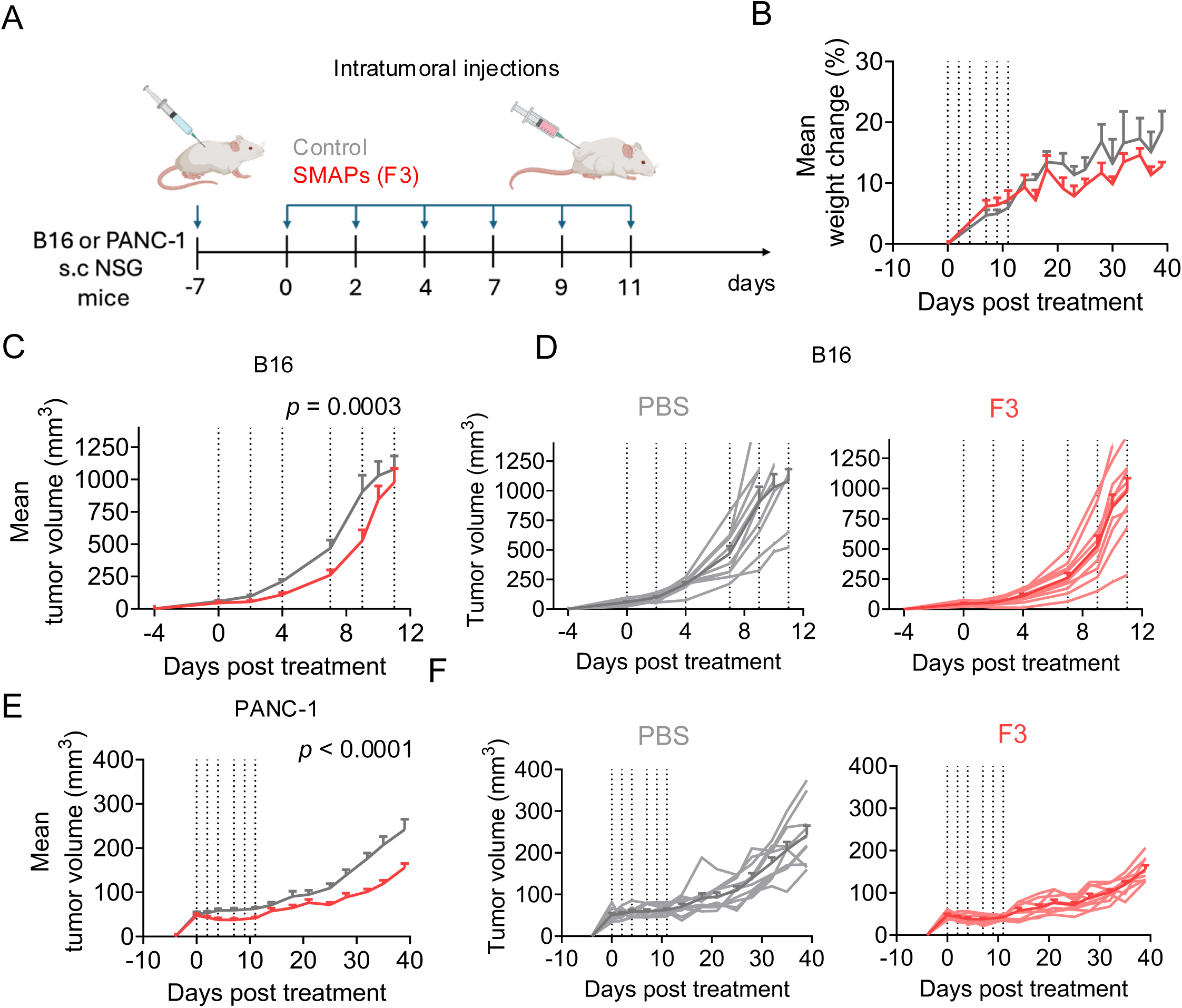
SMAPs (F3) control tumor progression in an aggressive mouse model of Melanoma B16 and slow growing PANC-1 Pancreatic cancer independent of lymphocytes. A) NSG mice bearing B16 murine melanoma or PANC-1 human pancreatic tumors were injected with 500μg/mouse SMAPS (F3) on days 0, 2, 4, 7, 9 and 11 intratumorally (i.t.). B) Weight changes relative to day 0 in PANC-1-bearing mice. C) Mean B16 tumor volume (mm^3^) measured with a digital calliper. D) Individual B16 tumor measurements post-treatment corresponding to C). E) Mean PANC-1 tumor volume (mm^3^). F) Individual PANC-1 tumor measurements post-treatment corresponding to E). Darker lines indicate the mean tumor volumes, and lighter lines indicate the tumor growth in individual mice. Data shown were pooled from 2 independent experiments. Data are presented as mean ± SEM. Statistical was performed using the 2way ANOVA with Tukey’s multiple comparisons test.

In conclusion, our study demonstrates the successful enrichment and characterization of SMAPs from NK92 cells using high-resolution size exclusion chromatography. The distinct protein compositions and functional properties of these NVEPs suggest their potential as novel therapeutic agents for solid tumours. It is possible that we were able to separate EV from SMAPs in NK-92 cell due to the deposition of a complement corona on the EVs, making them larger than most of the SMAPs. Further studies are needed to determine if the F2 associated complement and EV proteins are associated with the same particles. In vitro and ex vivo experiments confirmed a direct cytotoxic function of NK-92 SMAPs in suspension at readily achievable doses. In vivo experiments demonstrated that NK-92 SMAPs delay in vivo tumour growth in both aggressive B16 and slow growing PANC-1 tumour models. These findings highlight the therapeutic potential of SMAPs and provide a critical base-line for their future engineering.

## Discussion

Our study provides compelling evidence for the therapeutic potential of NVEPs derived from NK-92 cells, specifically SMAPs, in the treatment of solid tumours. The distinct isolation and characterization of these particles using high-resolution size exclusion chromatography (HR-SEC) allowed us to further delineate their unique protein compositions and functional properties.

The enrichment of F2 from the tandem SELC with complement-proteins requires further study. It is not clear if the human complement proteins are from 5% human serum used to grow the NK-92 cells or if this is made by the NK-92 cells. Such delivery of complement proteins by NK cells could play a role in enhancing the activity of antibodies, which are also an important effector arm for NK cells through CD16^18^.

On the other hand, SMAPs were enriched for proteins with direct cytotoxic potential, such as GZMB and PRF1. This is consistent with previous reports that SMAPs function as biological smart weapons, autonomously killing target cells (5). The ability of SMAPs to directly induce cytotoxicity without the need for prior sensitization makes them particularly attractive for targeting heterogeneous tumour cell populations that may evade immune detection. Our results further support the initial finding that THBS1 and THBS4 are associated with SMAPs. Our initial cryo-EM analysis didn’t provide direct insight into the nature of the THBS1 shell. Future correlative fluorescence cryo-structured illumination microscopy or direct labelling with antibodies linked to sign posts^19^ will be needed to address if any of the three structural sub-types: unbound, membrane bound or multilayer particles incorporate thrombospondins. The membrane bound F3 particles were ensheathed in 2-3 bilayer membranes, which is reminiscent of early observations of membrane bound particles containing cytotoxic proteins that were proposed to be guided by receptors from the cytotoxic lymphocyte through a membrane deposition process prior to release of cytotoxic granules^20^. This phenomenon may help to explain the presence of GZMB or PRF1 in CD81+ particles and the targeting of cytotoxic proteins by chimeric antigen receptors^21^. These particles are larger on average than other particles in F3, which are around 110 nm. This suggests that they may interact with the Sepharose resin to delay elution by a size independent mechanism. Alternative methods, such as iodixanol gradient centrifugation^16^, may be useful to further separate active components of F3.

The use of the NK-92 cell line, a GMP-grade cell line that has been used therapeutically, ensures the scalability and clinical applicability of our approach. The distinct isolation of SMAPs and their subsequent functional validation in an aggressive melanoma model provide a strong foundation for further development and optimization of SMAPs as therapeutic agents. EP preparations from NK-92 were previously shown to have activity in melanoma^22,23^. This activity was ascribed to exosomes, but it’s possible that SMAPs may have contributed to these effects, although we have shown that NK-92 cells release SMAPs only after prolonged high-density culture that would not be required for release of EVs.

Future studies should focus on optimising the production and purification processes for SMAPs from EVs to ensure consistent quality and yield. Additionally, exploring the engineering of these particles to enhance their specificity and efficacy against various tumour types could further improve their therapeutic potential. Investigating the mechanisms underlying the interactions between these particles and the tumour microenvironment will also provide valuable insights into their mode of action and potential combinatorial strategies with existing therapies. Preliminary studies suggest that many cell types produce thrombospondin-encapsulated dense particles. This may open the door to engineering non-cytotoxic cells to make cytotoxic SMAPs. A challenge may be that cytotoxic lymphocytes have evolved mechanisms to survive generation of such particles, including modifications to glycosphingolipid metabolism^24^.

In summary, our study establishes a novel approach for the isolation and characterization of SMAPs from NK92 cells, demonstrating their distinct therapeutic potential in solid tumour treatment. These findings pave the way for the development of engineered SMAP therapies, offering a promising new avenue for cancer treatment.

## Materials and Methods

### Cell Culture

The NK-92 cell line was maintained in complete RPMI-1640 medium (RPMI 1640 medium supplemented with 10% heat-inactivated fetal bovine serum, 10% human serum, 100μM non-essential amino acids, 10mM HEPES, 2mM L-glutamine, 1mM sodium pyruvate, 100U/mL of penicillin and 100μg/mL of streptomycin). PEG-EP were prepared from NK-92 supernatants using ExoQuick (Cambridge Biosciences) following the manufacturers protocol. Calu-3 cells were obtained from ATCC and were grown in Eagle’s Minimum Essential Medium with 10% FBS and other supplements as above. Lactate dehydrogenase (LDH) release was assayed to assess killing of Calu-3 cells (CyQuant, ThermoFisher).

#### CLL blood samples

Blood samples were obtained from untreated stage A patients. PBMCs were isolated on FicollPaque Gradient. Purification of CLL was performed using a Human B Cell Enrichment Kit II without CD43 Depletion (EasySep^TM^) according to manufacturer’s instructions.

#### Breast cancer tissue samples

Fresh breast cancer surgical specimens were dissociated enzymatically and mechanically using a tumor dissociation kit (Miltenyi Biotec) and gentleMACS™ Octo Dissociator. Negative selection of human tumor cells was performed using a tumor cell isolation kit (Miltenyi Biotec) according to manufacturer’s instructions.

EBV-transformed B cells (JY) were used as sensitive target cells.

Cells were cultured in RPMI 1640 GlutaMAX supplemented with 10% FCS and 50µmol/L 2-mercaptoethanol, 10mM HEPES, 1X MEM NEAA (Gibco), 1X Sodium pyruvate (Sigma), 10µg/ml ciprofloxacine (AppliChem).

Tissue samples and CLL blood samples have been obtained from the CRB Cancer of Toulouse, a structure recognized and approved by the French Ministry of Research (authorization N° DC-2020-4074). Approbation by the ethics department of the French Ministry of Research for the preparation and conservation of cell lines issued from cancer patients’ tissue samples has been obtained (authorization N° DC-2021-4673).

### Size-exclusion liquid chromatography (SELC)

The cells along with culture supernatant were transferred to 15 ml falcon tube and centrifuged at 300g for 10 mins. Supernatants were transferred to fresh 15 ml falcon tubes and centrifuged at 4,000g for 10 mins. The supernatant was concentrated using 100 kDa molecular weight cut-off Amicon centrifugal filter units (Merck Millipore) to 2 ml volume and loaded to Sepharose 4 Fast Flow column (10 mm x 300 mm; GE Healthcare) mounted to ÄKTA pure system (GE Healthcare) at a flow rate of 0.5 ml/min using PBS as the eluent. The absorbance chromatogram was recorded at 280 nm and 2 ml fractions were collected. The fractions corresponding to the void volume were pooled and concentrated using 10 kDa MW cut-off Amicon centrifugal filter units (Merck Millipore # UFC5010). This was followed by high resolution size exclusion chromatography (HR-SEC) by loading up to 5ml of the concentrated pooled particles to Sephacryl S-1000 Superfine HR (Cytiva, 17-0476-01) SELC column (x mm x y mm) connected to the ÄKTA Pure system with a flowrate of 0.5 mL/min using PBS as eluent. The absorbance chromatogram was recorded at 280 nm. The chromatograms showed 4 protein peaks and the corresponding fractions were pooled to generate F2 and F3 for further analysis. We note here that Sephacryl S-1000 Superfine HR was discontinued by Cytiva, but we have obtained similar results with Sephacryl S-500 Superfine HR (17-0613-01).

### Western blotting

Whole cell lysates (WCL) were prepared by lysing TH cells pellets in RIPA buffer (Thermo Fisher Scientific, #89901) containing Protease/Phosphatase inhibitor (PI) cocktail (CST; #5872) to a final concentration of 2 x 10^7^ cells/mL. The lysed cells were centrifuged at 10,000g for 10 mins at 4° C to remove cellular debris, and the protein supernatants were collected. The protein concentration was determined by BCA, protein samples were denatured in Laemmli buffer at 95° C for 10 mins and subsequently loaded on a SDS PAGE gel. In case of particles, isolated fractions were subjected to RIPA and PI treatment, the protein concentrations were measured using micro-BCA and equal amounts of proteins across fractions were loaded. Samples were resolved using 4-15% Mini-PROTEAN SDS-PAGE gels (Bio-Rad; #4561084), and transferred to 0.45 μm nitrocellulose membranes (Bio-Rad, #1620115), blocked with 5% BSA and incubated with the respective primary antibodies. Following incubation with primary antibodies, membranes were washed thrice with 1X TBST wash buffer and incubated with secondary antibodies. Primary and secondary antibody dilutions were used according to the manufacturer’s instructions. Membrane blocking with 5% BSA, or incubation with primary or secondary antibodies were performed either overnight at 4°C or 1 hr at room temperature. After incubation with secondary antibodies, membranes were washed four times with 1X TBST and image acquisition was performed using the Odyssey® CLx Near-Infrared detection system operated with Image Studio™ Lite quantification software (LICOR, Lincoln, NE).

### Nanoparticle tracking analysis (NTA)

NTA (Nanosight NS500, Malvern Instruments Ltd) with NTA 3.2 software set to light scattering mode was used to determine the concentration of particles. Videos for NTA were captured for 60 s, three sequential replicates per sample were obtained, and the three recordings were processed and averaged to determine the mean size and concentration of the particles. Particles were diluted in PBS to a concentration of 5E+8 to 1E+9 particles per ml for further analysis.

### Cryo EM

For Cryo-EM sample preparation, 3 µL of freshly concentrated sample were applied to glow-discharged Quantifoil R1.2/1.3 300 Cu mesh grids and vitrified using a Vitrobot Mark IV. Images were collected on a Titan Krios G4 equipped with a Falcon 4i camera and Selectris X energy filter at 64,000× magnification (1.9 Å/pixel). Particles were then manually segmented from the images using FIJI.

### Nano Flow Cytometry

The Nano flow cytometry analysis was performed using the Flow NanoAnalyzer (NanoFCM Co., LTD) according to the manufacturer’s instructions. Silica Nanospheres Cocktail (S16MExo, NanoFCM) was employed as the size standard to construct a calibration curve to convert side scatter intensities to particle size. NanoFCM 200 nm PS QC beads were used as a concentration standard for calculating particle concentration.

### Total Internal Reflection Fluorescent Microscopy (TIRFM)

TIRFM was performed on an Olympus IX83 inverted microscope equipped with a TIRF module. The instrument was equipped with an Olympus UApON 150 x 1.45 NA oil-immersion objective and Photomertrics Evolve delta EMCCD camera. Image analysis and visualization were performed using ImageJ (National Institutes of Health)^12^. For TIRFM, the particles were labelled with PRF1 (Biolegend #308108, AF488), GZMB (Biolegend #372219 AF647) and CD81 (Santa Cruz, sc-166029 AF546) antibodies or WGA (Biotium, Fremont, CA; 29027-1 CF405S) and captured onto poly-L lysine coated glass^13^. For dSTORM imaging, particles were mounted with a reducing buffer system. A total of 10,000 images were captured on a Olympus MITICO TIRFM microscope with a 150x TIRF objective and an EM-CCD camera or a Nanoimager (Oxford Nanoimaging) with a 100x oil-objective lens, and captured images were analyzed with Fiji or Nanoimager Software v1.4.8740 (Oxford Nanoimaging). For the particle segmentation analysis Fig S1B, a mask was created from each channel to detect individual particles using a local thresholding method and Gaussian Blur filters in Fiji (ImageJ). Percentages of single, double and triple positive particles were calculated from the total number of particles detected.

### Multicolor Flow Cytometry (FCM)

The staining was performed in 5% BSA in PBS pH 7.4 (0.22 µm-filtered) at +4°C for at least 30 mins. Further, cells were washed thrice and then acquired using an LSRFortessa X-20 flow cytometer with a High Throughput Sampler (HTS). To ensure absolute quantification, Quantum Molecules of Equivalent Soluble Fluorescent dye (MESF) beads were used to set photomultiplier voltages to position all the calibration peaks within an optimal arbitrary fluorescence units’ dynamic range (between 10^1^ and 2 × 10^5^, and before compensation). Fluorescence spectral overlap compensation was then performed using single colour-labelled cells and unlabelled cells. To calculate compensation matrixes for markers with low surface expression levels, unstained and single colour stained UltraComp eBeads were used. After applying the compensation matrix, experimental specimens and Quantum MESF beads were acquired using the same instrument settings.

### Proteomics

Particles (1 ml) were loaded onto a pre-washed 10 kDa MW cut-off spin filter (Vivacon 500, Sartorius) and centrifuged at 14,300 rcf for 10 min. Protein was denatured on-filter in 8 M urea, 100 mM triethylammonium bicarbonate pH 8.5 for 30 min at ambient temperature. Cysteine reduction and alkylation were performed for 30 min at ambient temperature with 10 mM tris(2-carboxyethyl)phosphine and 50 mM iodoacetamide, respectively. The samples were centrifuged as before then washed twice with 50 mM triethylammonium bicarbonate pH 8.5. Digestion was performed overnight at 37°C with 2 µg trypsin (sequencing grade, Promega) in 50 mM triethylammonium bicarbonate pH 8.5, then peptides were eluted by centrifugation of the digest buffer, then 0.1% TFA, then 50% acetonitrile 0.1% TFA. Peptides were dried by vacuum centrifugation then reconstituted in 5% formic acid, 5% DMSO immediately prior to LC-MS analysis. Peptides were separated using a Dionex Ultimate 3000 UPLC (Thermo Fisher Scientific) and analysed using a Q Exactive mass spectrometer (Thermo Fisher Scientific). First, peptides were loaded onto a trap column (PepMapC18; 300 µm x 5 mm, 5 µm particle size, Thermo Fischer Scientific) for 1 min at 20 μL/min flowrate followed by chromatographic separation on a 50 cm-long EasySpray column (ES803, Thermo Fischer Scientific) with a gradient of 2 to 35% acetonitrile in 0.1% formic acid and 5% DMSO at 250 nL/min flow rate for 60 min. The Q Exactive was operated in between full MS scan and MS/MS acquisition. Survey scans were acquired in the Orbitrap mass analyser over an m/z window of 380-1500 and at a resolution of 70k (AGC target at 3e6 ions). Prior to MS/MS acquisition, the top fifteen most intense precursor ions (charge state >=2) were sequentially isolated in the Quad (m/z 1.6 window) and fragmented on the HCD cell (normalised collision energy of 28). MS/MS data were obtained in the orbitrap at a resolution of 17.5K with a maximum acquisition time of 128 ms, an AGC target of 1e5 and a dynamic exclusion of 27 seconds. For proteomic analyses, the raw files were searched against the reviewed Uniprot Homo sapiens database (retrieved 2,01,80,131) using MaxQuant v1.6.10.43. The LFQ intensities were normalized using particle concentration from NTA and further average-based normalization was carried out to obtain the enriched set of proteins in each fractions.

### Cell viability assay

Cells were seeded in 96 well tissue culture plates at a density of 10,000 cells per well, and after 24hrs of seeding, the cell culture medium was removed. This was followed by a wash with PBS. The particles were diluted in cell culture medium at different particle concentrations (particles: 1E6, 5E6, 1E7, 5E7, 1E8 and 5E8). In the control group only 100uL culture medium was added while the experimental group cells were treated with particles. The experimental group included particles dilutions from NK92 cells. Cells were left in cell culture incubator for 48 hrs. Following incubation, 100µL fresh media with 10µL MTT reagent (Roche # 11465007001) was added to the cells and they were incubated for 4 hrs at 37°C. After incubation, media with MTT reagent was removed from wells, cells were lysed and formazan crystals were solubilized with a lysis buffer (Roche # 11465007001). Using microplate readers, the absorbance of formazan solutions was recorded at 570 nm with background correction at 750nm. Cell viability and cell death was calculated as percentage of controls cells as % live cells = (absorbance of cells treatment group/absorbance of control cells) *100.

In some experiments 1 x 10^4^ JY cells, 1 x 10^4^ primary breast cancer cells and 1 x 10^5^ CLL cells were seeded in 96-well flat bottom plates. Cells were treated with different concentrations of SMAPs (F3 Fraction) for 6h or 24h (JY cells) and 24h (CLL and primary breast cancer cells) at 37°C, 5% CO2. Cells were transferred in U-bottom 96-well plates and stained with the viability dye eFluor™ 780 (eBioscience™ #65-0865-14) to be analyzed by flow cytometry. Acquisitions were done with a MACSQuant Analyzer 10 flow cytometer (Miltenyi Biotec) and analyzed using FlowJo 10.

### In vivo experiments

The NOD scid gamma (NSG) female mice (7-9 weeks of age) were purchased from Charles River, UK. Mice were anesthetized with isoflurane prior to subcutaneous injection on the right flank of 0.5E+6 B16F10-OVA or PANC-1 cells/100μL/mouse with a 23G needle. The B16F10-OVA cells were a gift from the Yang Shi lab, University of Oxford, UK. PANC-1 cells were injected with Matrigel (cat. 3632-010-02), 2:1 cells/Matrigel volume ratio. Seven days later, when the tumors were palpable, mice received 500μg/30μL/mouse of SMAPS (F3) or vehicle control (PBS) intratumorally (i.t.) with a 29G insulin syringe. The i.t. injections were repeated 3 times a week for 2 weeks (days 0, 2, 4, 7, 9 and 11). The tumors were measured 3 to 4 times per week using a digital calliper, and the tumor volume (V) was calculated with the formula: V = [[[width2 (mm)] x length (mm)] x 0.52]. Mice reached the humane endpoint when the tumors reached 500-1000 mm3. Ethical approval of animal experiments was obtained under the license PPL PP8978826. Data visualization and statistical analysis were performed on GraphPad Prism version 10.

### Statistical analysis

Statistical analysis was conducted utilizing GraphPad Prism software v10.0.3. To assess statistical significance, various methods were employed, including normality tests like Kolmogorov-Smirnov or Shapiro-Wilk tests, followed by appropriate tests such as Mann-Whitney test, unpaired t-test, or multiple unpaired t-tests.

## Acknowledgments

We thank S. Balint for initial analysis of SMAPs released from NK-92 cells. JC was supported by a Cancer Research Institute Irvington Fellowship (CRI4503).

## Figure, video and table legends

**Supp Fig 1.**
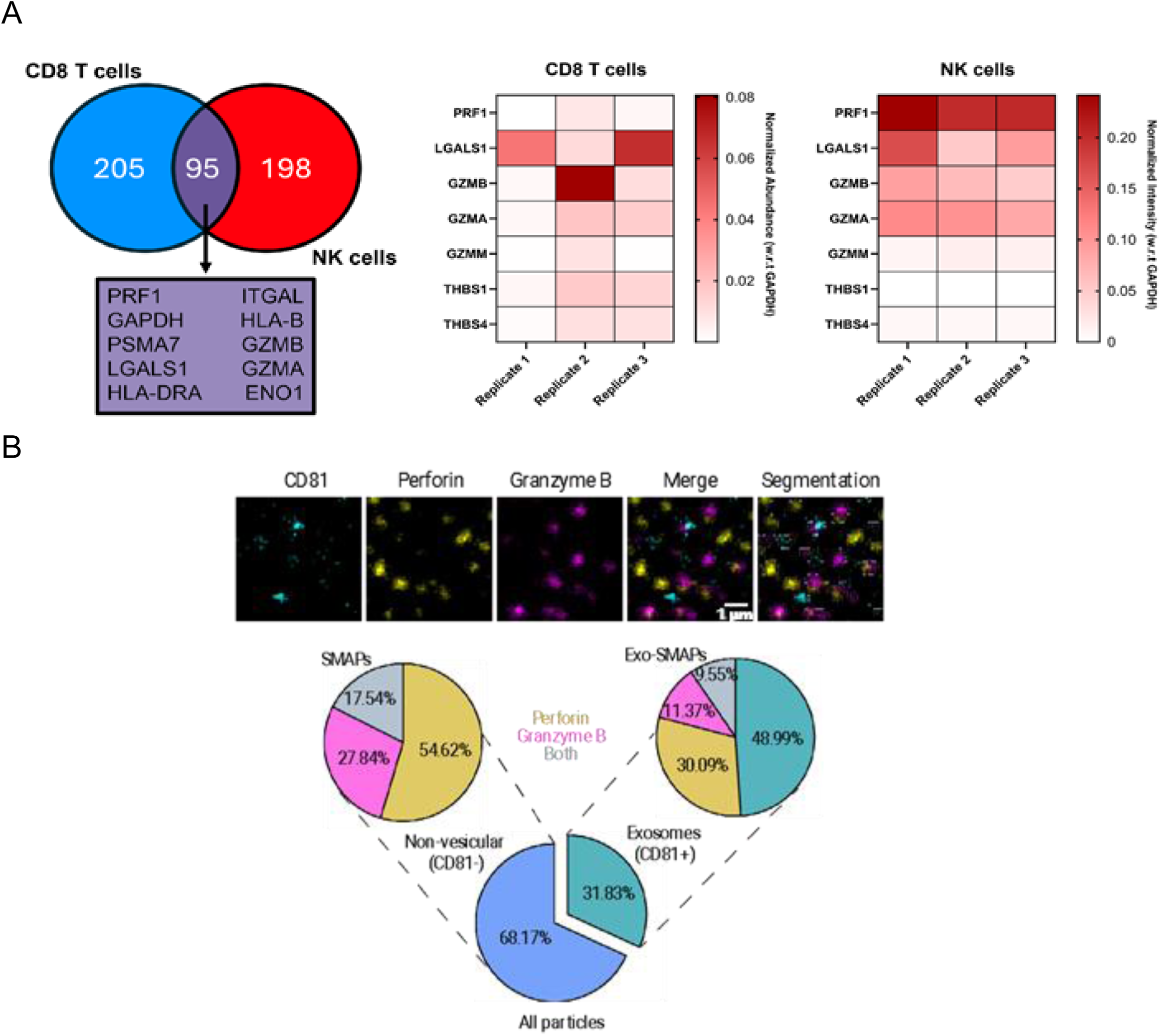
4FF derived NK92 Extracellular particles produce EVs and SMAPs. (A) Quantitative proteomics of 4FF derived NK92 Extracellular Particles (EP) and comparison to shared and SMAP signature from Balint et al., 2020 (B) NK92 EP from 4FF were adsorbed on to glass slides and immunostained for CD81 (cyan), PRF1 (yellow) and GZMB (magenta), and imaged with TIRFM. CD81 positive particles were segmented to quantify the proportions of exosomes and CD81 negative particles were quantified under non-vesicular particles. Data shows the pie chart distribution of particles positive for exosomes (green), GZMB (pink), PRF1 (yellow) and double positive for GZMB and PRF1 (grey). Quantification is based on a total of 46052 particles from 25 fields of view.

**Supp Fig 2.**
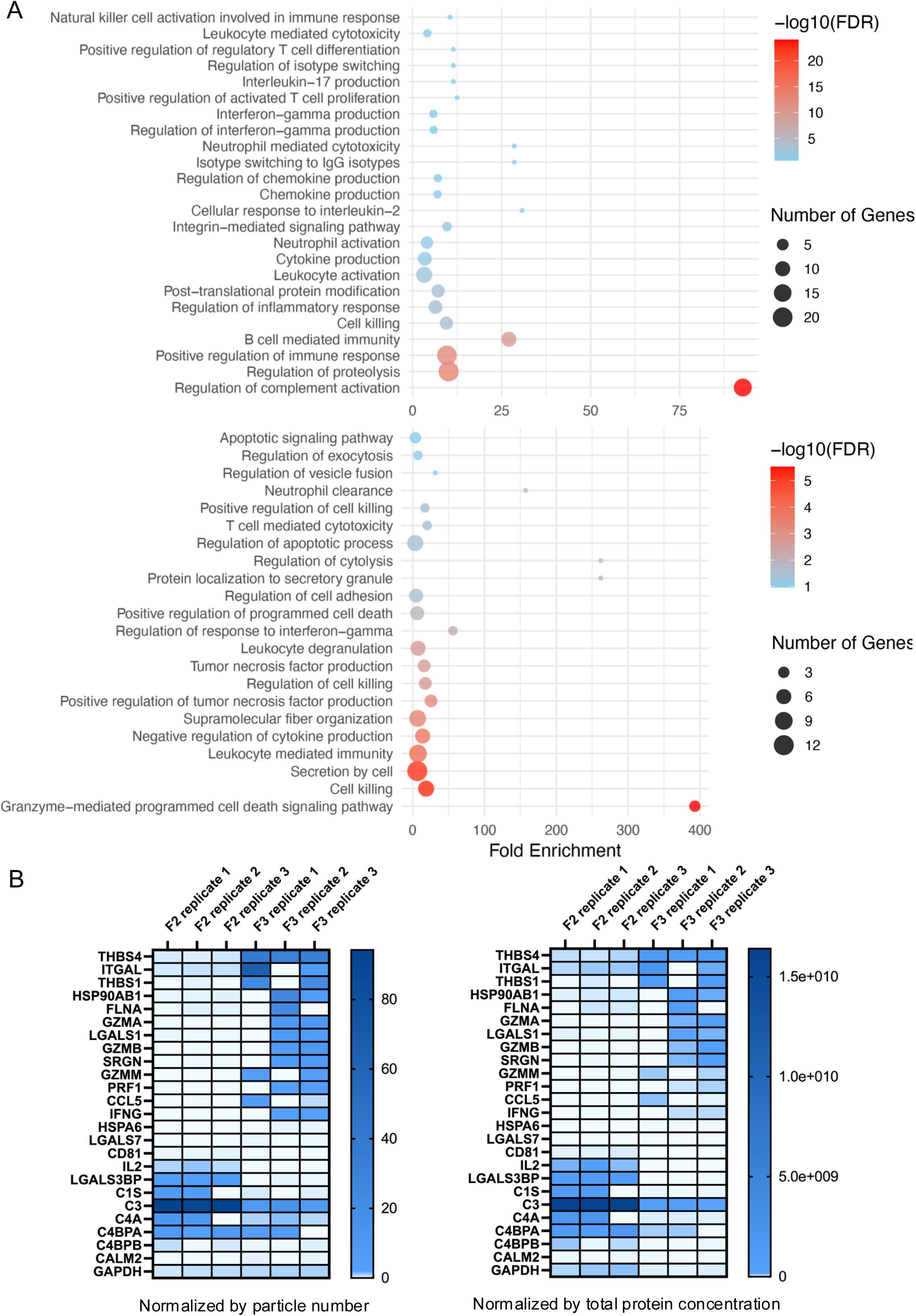
SMAPs (F3) fraction is enriched for Granzyme mediated apoptosis associated proteins and ComEV (F2) are enriched for complement activation associated proteins and the trend is consistent across different normalization methods. (A) Pathway analysis of enriched proteins from F2 and F3 fractions is represented using a bubble plot. (B) Particle and protein concentration normalized intensities of SMAP and EV signature proteins along with complement proteins plotted across proteomics replicates in relation to Fig 2B

**Supp Video 1 SMAPs (F3) drives caspase driven cell death in NALM-6 cells associated with Figure 3D-E**

**Supp Fig 3.**
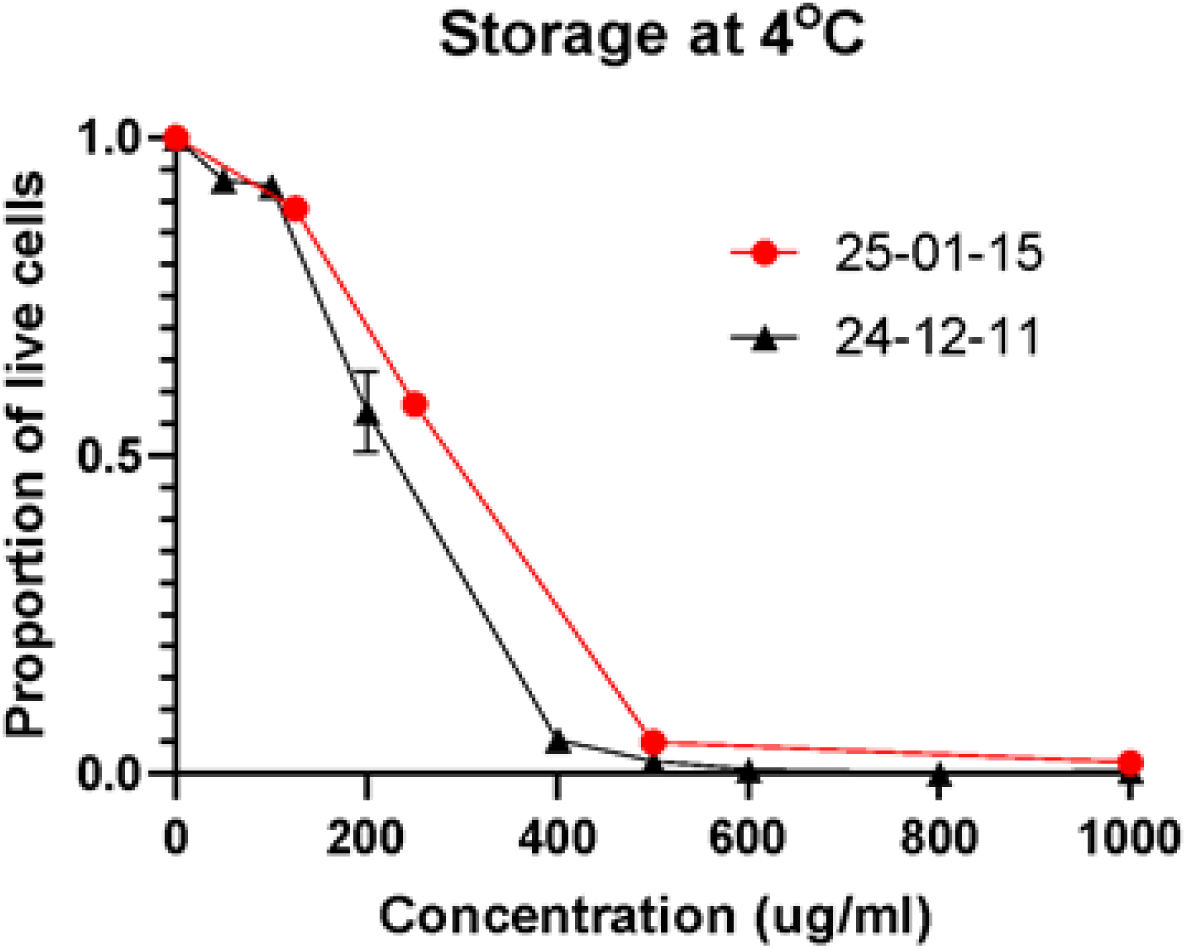
Ca2+ stabilized SMAPs are functional if kept at at 4° C for a month.

**Supp Table 1.** Table of protein concentration as measured by BCA, across different stages of particle purification using SELC.

**Figure 5**
|  |  | BCA repeat 1 | BCA repeat 2 | BCA repeat 3 |
| --- | --- | --- | --- | --- |
| Crude supernatant |  | 16 mg/ml | 25 mg/ml | 17 mg/ml |
| Sephrose column - SELC | Pooled particles | 10 mg/ml | 17 mg/ml | 12 mg/ml |
| S1000 high resolution- SELC | F2 (EV enriched) | 25 mg/ml | 33 mg/ml | 23 mg/ml |
|  | F3 (SMAP enriched) | 30 mg/ml | 41 mg/ml | 31 mg/ml |

